# Repopulating Kupffer Cells Originate Directly from Hematopoietic Stem Cells

**DOI:** 10.1101/2020.07.31.230649

**Authors:** Xu Fan, Pei Lu, Xianghua Cui, Peng Wu, Weiran Lin, Dong Zhang, Shongzong Yuan, Bing Liu, Fangyan Chen, Hong You, Handong Wei, Fuchu He, Jidong Jia, Ying Jiang

**Author notes:** **Corresponding Authors: Ying Jiang**, State Key Laboratory of Proteomics, Beijing Proteeome Research Center, National Center for Protein Sciences (Beijing), Beijing Institute of Lifeomics, Beijing 102206, China., **Jidong Jia**, Liver Research Center, Beijing Friendship Hospital, Capital Medical University, Beijing Key Laboratory of Translational Medicine in Liver Cirrhosis & National Clinical Research Center of Digestive Diseases, Beijing 100050, China.; **Fuchu He**, State Key Laboratory of Proteomics, Beijing Proteeome Research Center, National Center for Protein Sciences (Beijing), Beijing Institute of Lifeomics, Beijing 102206, China.

## Abstract

Kupffer cells (KCs) originate from yolk sac progenitors before birth. Throughout adulthood, they self-maintain independently from the input of circulating monocytes (MOs) at stead state, and are replenished within 2 weeks after having been depleted, but the origin of repopulating KCs in adult remains unclear. The current paradigm dictates that repopulating KCs originate from preexisting KCs or monocytes, but there remains a lack of fate-mapping evidence. In current study, we firstly traced the fate of preexisting KCs and that of monocytic cells with tissue-resident macrophage-specific and monocytic cell-specific fate mapping mouse models, respectively, and found no evidences that repopulating KCs originate from preexisting KCs or MOs. Secondly, we performed genetic lineage tracing to determine the type of progenitor cells involved in response to KC depletion in mice, and found that in response to KC depletion, hematopoietic stem cells (HSCs) proliferated in the bone marrow, mobilized into the blood, adoptively transferred into the liver and differentiated into KCs. Finally, we traced the fate of HSCs in a HSC-specific fate-mapping mouse model, in context of chronic liver inflammation induced by repeated carbon tetrachloride treatment, and confirmed that repopulating KCs originated directly from HSCs. Taken together, these findings provided strong in vivo fate-mapping evidences that repopulating KCs originate directly from Hematopoietic stem cells not from preexisting KCs or from MOs.

**Significance:** There is a standing controversy in the field regarding the cellular origin of repopulating macrophages. This paper provides strong in vivo fate-mapping evidences that repopulating KCs originate directly from hematopoietic stem cells not from preexisting KCs or from MOs, which presenting a completely novel understanding of the cellular origin of repopulating Kupffer Cells and shedding light on the divergent roles of KCs in liver homeostasis and diseases.

## Introduction

Kupffer cells (KCs), the tissue resident macrophages (TRMs) in the liver, play crucial roles in liver homeostasis and in the pathogenesis of liver diseases^1^. According to the common mononuclear phagocyte system theory, all TRMs including KCs originate from and are continuously replenished by circulating MOs^2^. However, the concept has been being undermined by the new insight that the majority of TRMs including KCs originate from yolk sac Erythro-Myeloid Progenitors^3^. Furthermore, unlike skin and intestine, adult liver resists the colonization of monocyte-derived macrophages, and retains fetal-derived KCs with potential of long-term self-maintaining^4,5^, at steady state.

Previous studies showed KCs are replenished within two weeks even following a severe depletion^6^. However, the cellular origin of repopulating KCs remains unclear. It has been suggested that preexisting KCs^3,7^ or MOs^8,9^ were the cellular origin of repopulating KCs. The former hypothesis is mainly supported by the findings that repopulation of KCs is independent of the signal of CC chemokine receptor 2 (CCR2), a chemokine receptor predominantly expressed on monocytes^5^, that KCs have the potential of proliferation in vitro upon inactivation of transcription factors MafB and c-Maf^10^, and that KCs proliferate actively in context of glucan-induced granuloma formation^11^. The second hypothesis is mainly supported by the findings that a partial replacement of KCs by bone marrow (BM)-derived progenitors is observed in BM transplantation experiments^12^, in adoptive transfer experiments^9^, and in severe experimental Listeria infection^13^. However, both of these hypotheses have yet to be confirmed by in vivo fate mapping evidence.

Therefore, the purpose of current study was to determine whether repopulating KCs originate from preexisting KCs or from MOs, as previously reported, using genetic inducible fate mapping and, if not, to determine what type of progenitor cells give rise to repopulating KCs. For this purpose, we firstly tracing the fate of preexisting KCs and that of MOs during KC-repopulation, in a TRM- and a monocyte-specific genetic inducible fate mapping mouse model, we found no evidences that repopulating KCs originate from preexisting KCs or from MOs. Then, using genetic lineage tracing we found that hematopoietic stem cells (HSCs) act as progenitor cells in response to KC-depletion. Finally, employing a HSC-specific fate-mapping system, we confirmed that repopulating KCs originate directly from HSCs, in context of chronic liver inflammation induced by repeated carbon tetrachloride (CCl_4_) treatment, a common used mouse model associated with KC-depletion without MO-depletion.

## Results

### Repopulating KCs did not originate from preexisting KCs

Given that adult mouse TRMs originate from colony-stimulating factor 1 receptor (Csf1r)-expressing yolk-sac progenitors^14^, an inducible Csf1r^MeriCreMer^ fate-mapping system is wildly used for labeling Csf1r-expressing yolk-sac progenitors and to follow their progeny in adult mice^15^. Accordingly, we traced the fate of preexisting KCs during KC repopulation as follows.

To label KCs, Csf1r^CreERT2^ activity was induced with a pulse of tamoxifen in Csf1r^MeriCreMer^; Rosa^mT/mG^ mouse embryos at E8.5. To selectively deplete KCs without depleting bone marrow macrophages (Fig. S1 *A* and *B*), and without triggering liver inflammation (Fig. S1 *C* and *D*), 20mg/kg of Clo was intraperitoneally injected into pulsed mice 8 weeks after birth. To determine the contribution of “non-KCs” to KC repopulation, we compared the KC-labeling index before and after KC repopulation. The rationale for this approach was as follows: if repopulating KCs originate from genetically labeled preexisting KCs, then the KC-labeling index should remain unchanged. In contrast, if repopulating KCs originate from unlabeled progenitor cells, then the KC-labeling index should decrease^16^.

A tamoxifen pulse at E8.5 resulted in labeling that was completely restricted to KCs and did not extend to HSCs or blood leukocytes. No labeled KCs were detected in tamoxifen-treated Csf1r^wt^; Rosa^mT/mG^ animals (Fig. 1*A*). Furthermore, no differences were observed in the labeling indexes of the MHC-II^+^ or MHC-II^−^ KC subgroups ^17^or the CD68^+^ or CD68^−^ KC subgroups^18^ (Fig. 1*B* and Fig. S2*A*), indicating that the labeling was not restricted to a specific KC subgroup.

**Fig. 1.**
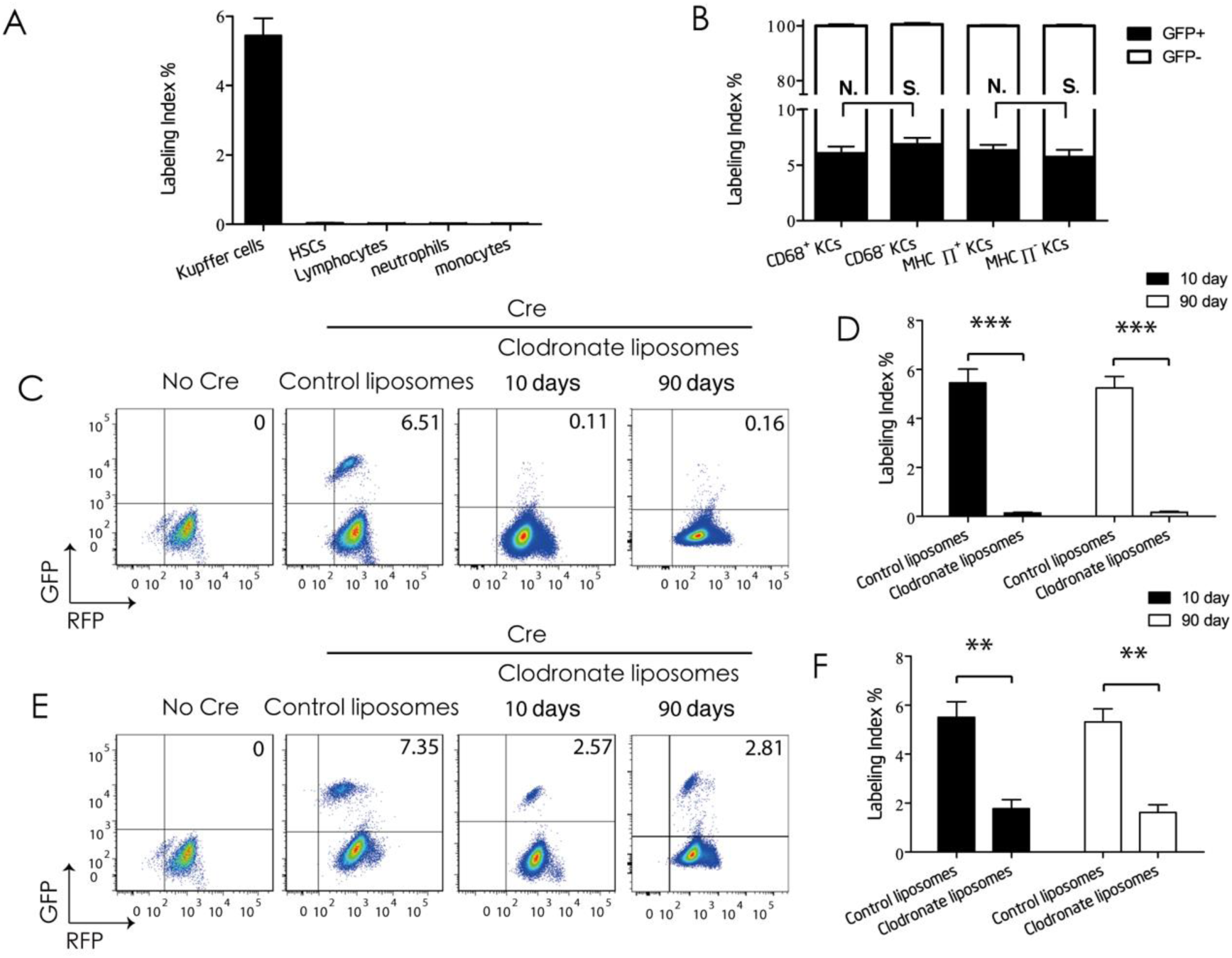
Repopulating Kupffer cells do not originate from preexisting Kupffer cells. (*A*) Label index of Kupffer cells (**KCs**), hematopoietic stem cells (**HSCs**), lymphocytes, neutrophils, and monocytes **(MOs)** from E8.5-pulsed Cx3cr1^CreERT2^; Rosa^mT/mG^ mice (**Cre mice**). Values are the means ± SEM from 6 samples. (*B*) Label index of CD68^+^ or CD68^−^, and MHCII^+^ or MHCII^−^ KC subgroups from Cre mice. Values are the means ± SEM from 6 samples. **N.S.** No significant difference between indicated groups by *t*-test. (*C*) Flow cytometric analysis of KCs from E8.5-pulsed Csf1r^Wt^; Rosa^mT/mG^ mice (**No Cre mice**) or from Cre mice at 10 day and 90 day post-intraperitoneal injection (**i.p.**) with 20mg/kg of control-liposomes or 20 mg/kg of clodronate-liposomes. (*D*) Label index of KCs from indicated mice analyzed in *C*. Values are the means ± SEM from 6 samples. *** *P* < 0.001 between groups by *t*-test. (*E*) Flow cytometric analysis of KCs from No Cre mice or from Cre mice at 10 day and 90 day post i.p. with 10 mg/kg of control-liposomes or 10mg/kg of clodronate-liposomes. (*F*) Label index of KCs from indicated mice analyzed in *E*. Values are the means ± SEM from 6 samples. ** *P* < 0.01 between groups by t-test.

After complete repopulation following 90% KC depletion (10 to 90 days post Clo injection), the mean KC-labeling index for the clodronate-liposomes group (repopulation after depletion) was reduced by approximately 95% compared with the control-liposomes group (no depletion) [0.14 ± 0.06 vs. 5.42 ± 1.44%] (Fig. 1 *C* and *D*). The proliferative ability of KCs might be impaired because of their near-complete depletion. To exclude this possibility, only 60% of KCs were depleted by intraperitoneal injection of 10 mg/kg clodronate-liposomes in 8-week-old Csf1r^MeriCreMer^; Rosa^mT/mG^ mice (Fig. S2 *B* and *C*), pulsed with tamoxifen at E8.5. Similarly, the mean KC-labeling index for the clodronate-liposomes group was reduced by 70% compared with the control-liposomes group [1.79 ± 0.69 vs. 5.41 ± 1.62%] (Fig. 1 *E* and *F*). Moreover, we found that the KC labeling index remained unchanged through 90 day post KC repopulation (Fig. S2 *D* and *E*).

Together, these results demonstrated that repopulating KCs originate from unlabeled progenitor cells rather than genetically labeled preexisting KCs.

### Repopulating KCs originated from hematopoietic progenitors in bone marrow

Next, we attempted to determine what type of progenitor cells give rise to repopulating KCs. Previous studies demonstrate that some KCs are of donor hematopoietic progenitor origin in BM chimeras^12^. However, such transplantation protocols do not accurately reflect KC repopulation under physiological conditions. In particular, that protocol involved total-body irradiation, which could affect peripheral cell entry into the liver by impairing the integrity of the hepatic sinusoid^7^. Therefore, we traced the fate of hematopoietic progenitors during KC repopulation under physiological conditions as follows. Purified HSCs (defined as Lin^neg^Sca-1^+^c-kit^+^CD34^−^CD135^−^CD48^−^CD150^+^) from B6GFP transgenic mice were engrafted into Kit^w^/Kit^wv^ recipients, which can accept HSC grafts in the C57BL/6 background without myeloablation^19^.

To selectively deplete KCs by approximately 90% or 60%, 20mg/kg or 10mg/kg of Clo were intraperitoneally injected into HSCs chimeras, respectively, at 8 weeks post-engraftment. To determine whether hematopoietic progenitors contribute to KC repopulation, the total number of host-origin KCs after depletion and after complete repopulation were compared. The rationale for this approach was that if repopulating KCs originate from donor-origin hematopoietic progenitors then the total number of host-origin KCs should remain unchanged throughout repopulation.

We found that 8 weeks after engraftment, only hematopoietic cells (including HSCs, MOs, neutrophils, and most lymphocytes) not KCs within the recipients were of donor HSC origin. (Fig. 2 *A, B* and Fig. S3). For the observation period from 10 days till 90 days post Clo injection (0 day and 80 days post complete KC repopulation), all repopulating KCs were labeled with GFP (Fig. 2 *C-F*). These results demonstrate that all repopulating KCs originate from hematopoietic progenitors in the BM.

**Fig. 2.**
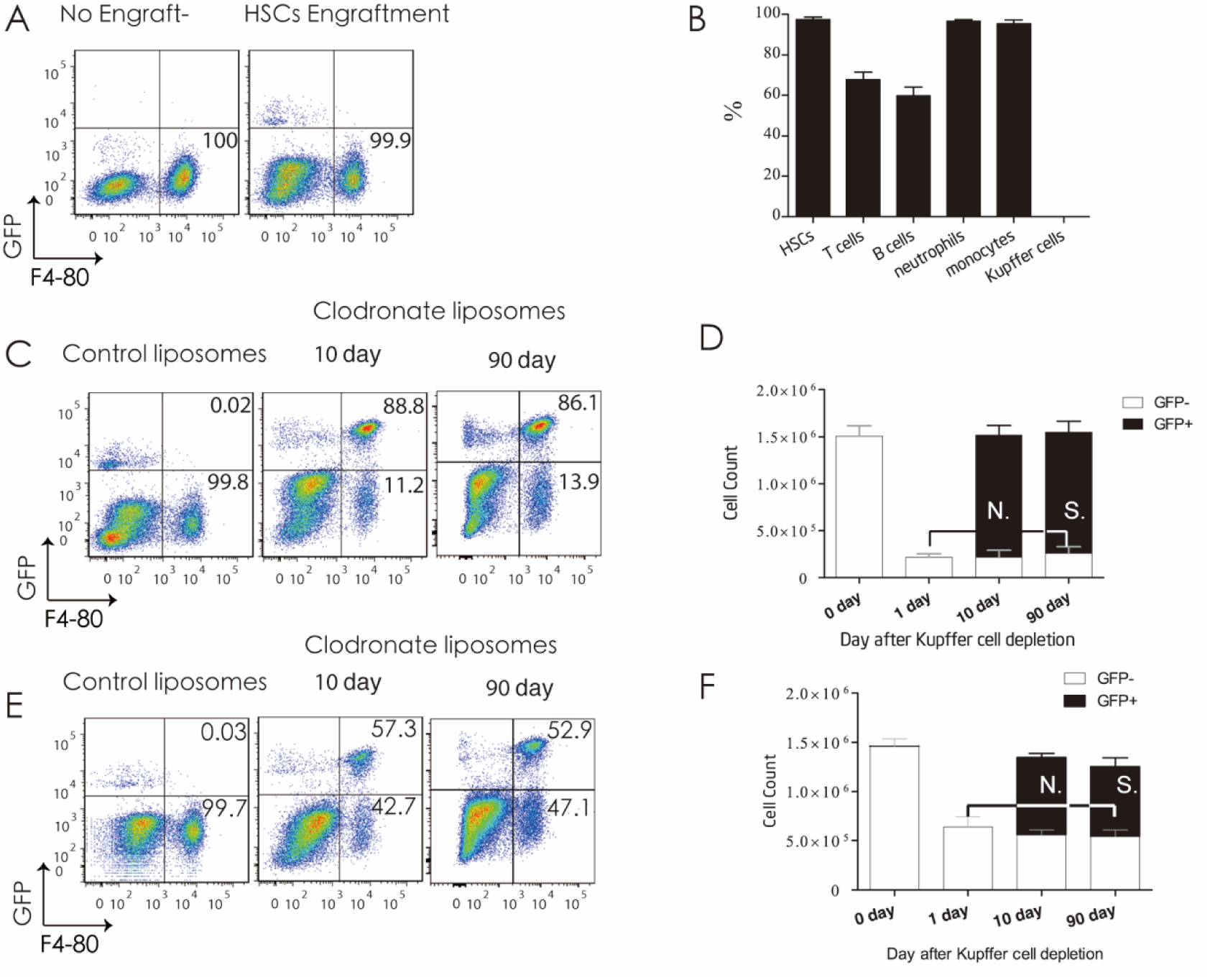
Repopulating Kupffer cells originate from hematopoietic progenitors. (*A*) Flow cytometric analysis of liver non-parenchymal cells from purified GFP^+^ HSC-chimeric Kit^w^/Kit^wv^ mice before engraftment (**no engraftment**) and 8 week post-engraftment (**HSC-engraftment**). (*B*) Donor chimerism of HSCs, T-cells, B-cells, neutrophils, MOs, and KCs from HSC-chimeric Kit^w^/Kit^wv^ mice analyzed in A. Values are the means ± SEM from 3 samples. (*C*) Flow cytometric analysis of liver non-parenchymal cells from HSC-chimeric Kit^w^/Kit^wv^ mice at 10 and 90 day post-i.p. with 20mg/kg of clodronate-liposomes (**90% KC-depletion**) (*D*) Cell counts of KCs from HSC-chimeric Kit^w^/Kit^wv^ mice at 0, 1, 10, and 90 day after 90% KC depletion. Values are the means ± SEM from 4 samples **N.S.**, no significant difference between groups by ANOVA. (*E*) Flow cytometric analysis of liver non-parenchymal cells from HSC-chimeric Kit^w^/Kit^wv^ mice at 10 and 90 day post-i.p. with 10 mg/kg of clodronate-liposomes (**60% KC-depletion**). (*F*) Cell counts of KCs from HSC-chimeric Kit^w^/Kit^wv^ mice at 0, 1,10 and 90 day after 60% KC depletion. Values are the means ± SEM from 4 samples **N.S.**, no significant difference between groups by ANOVA.

### Repopulating KCs did not originate from MOs

We then sought to test the hypothesis that repopulating KCs originate from MOs. For this purpose, we induced Cre activity by Tamoxifen administration in adult Cx3cr1^CreERT2^; Rosa^YFP^ mice^20^, and then dynamically investigated the labeling index of monocytic cells. We found that during an observation period of 2 to 25 days after 5 days of consecutive tamoxifen administration, Cx3cr1^CreERT2^; Rosa^YFP^ mice showed labeling in all monocytic cells, including macrophages and DC progenitors (MDPs), common monocyte progenitors (cMoPs), monocytes in BM, Ly6C^hi^ or Ly6C^low^ monocytes in the blood (Fig. S4*A*), and intra-splenic MOs (Fig. S4 *B* and *C*). However, no labeled cells were detected in HSCs or KCs (Fig. S4*D*). As expected, no labeled monocytic cells were detected in tamoxifen-treated Cx3cr1^wt^; Rosa^YFP^ animals (Fig. S4*E*).

Thus, we traced the fate of circulating MOs and of intra-splenic MOs during KC repopulation as follows. To label monocytic cells, adult Cx3cr1^CreERT2^; Rosa^YFP^ mice were pulsed with 5 days of consecutive of tamoxifen administration. A 15-day wash-out period was conducted to allow tamoxifen levels to dissipate before the initiation of KC depletion^21^. To determine whether MOs contributed to KC repopulation, the labeling-index ratio of KCs to both Ly6C^hi^ and Ly6C^low^ circulating monocytes, and the labeling index of KCs to intra-splenic MOs were compared to a constant value (0.9) at 10 days post-clodronate-liposomes injection.

The rationale for this approach was as follows: following complete repopulation after 90% KC-depletion, if repopulating KCs originate from either circulating MOs or intra-splenic MOs, then the labeling index ratio of repopulating KCs vs. Ly6C^hi or low^ MOs should be close to 0.9. However, if repopulating KCs do not originate from MOs and do not express the Cx3cr1 promoter during differentiation, then the KC labeling-index should be close to zero. Finally, if repopulating KCs do not originate from MOs but do express from the Cx3cr1 promoter during differentiation, and if low levels of Cre activity persist for 3 half-lives^22^ (15 days) after tamoxifen administration, then both ratios should be less than 0.9.

We found that throughout KC repopulation (from 4 to 10 days post Clo injection),both the label index ratio of KCs vs. circulating MOs and the labeling index ratio of KCs vs intra-splenic MOs were less than 0.9 (Fig. 3, Fig. S4 *F - I*). At 45 days and 90 days post Clo injection, although no labeled monocytic cells were detected (Fig. S4*A*), the labeling index of repopulating KCs remained unchanged (Fig. 3*B*). Taken together, these results demonstrate that repopulating KCs originate from unlabeled non-monocytic hematopoietic progenitors rather than labeled MOs.

**Fig. 3.**
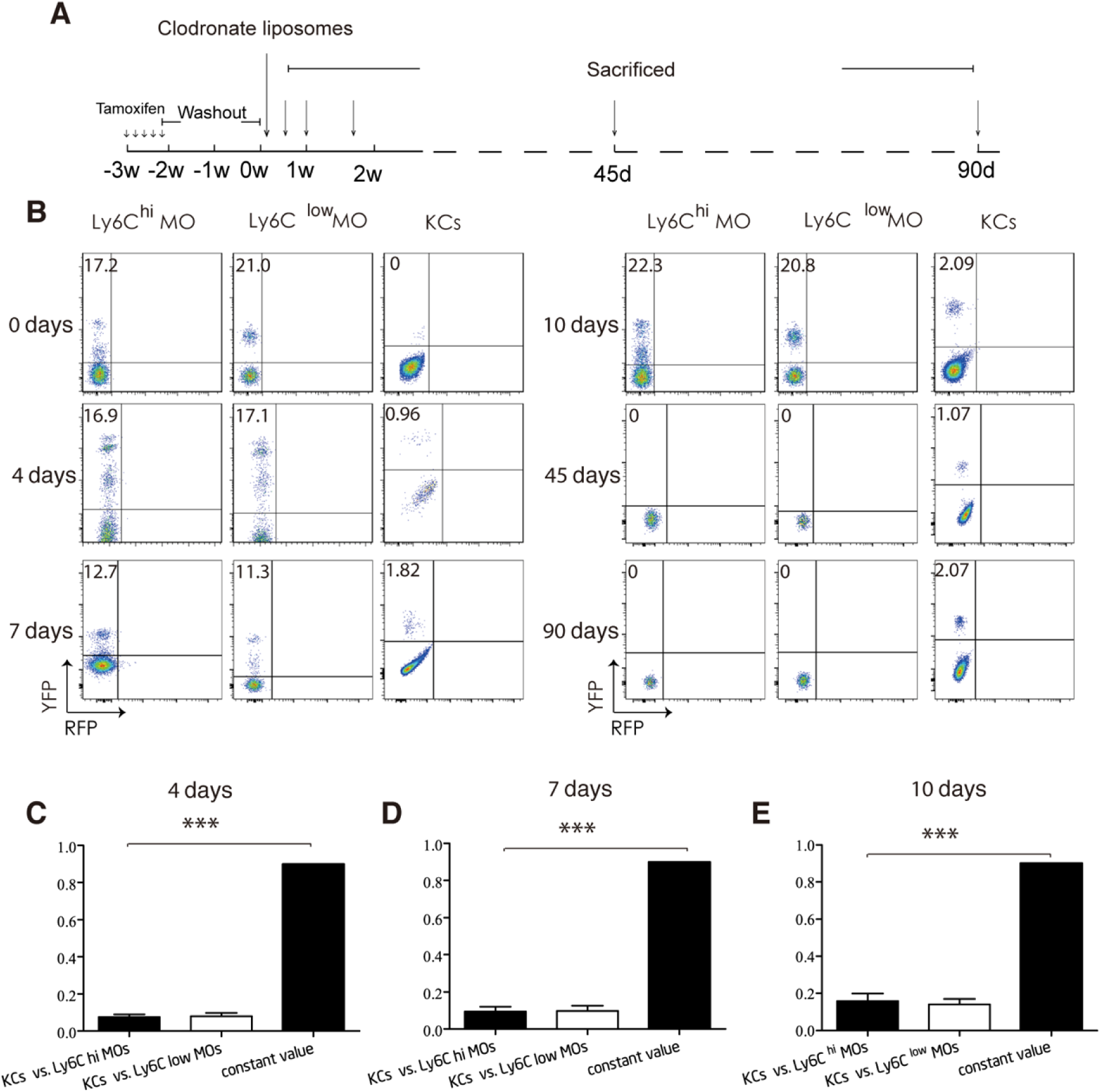
Repopulating Kupffer cells do not originate from MOs. (*A*) Experimental schedule (*B*) Flow cytometric analysis of indicated cells from adult-pulsed Cx3cr1^CreERT2^; RosaYFP mice (Cre mice) i.p. with 20mg/ml clodronate-liposomes (KC-depletion) at indicated time point post injection (n = 5/group). (*C*) (*D*) (*E*) Label index of indicated cells from Cre mice at indicated time point post-i.p. with 20mg/kg of control-liposomes analyzed in *B*. Values are the means ± SEM from 6 samples. \*\*\**P* < 0.001 between groups by ANOVA.

### Hematopoietic stem cells act as progenitors in response to Kupffer cell depletion

Next, we sought to investigate which type of non-monocytic hematopoietic progenitors give rise to repopulating KCs. According to previous reports^23^, the progenitor clls for repopulating KCs should have a context-dependent probability of differentiating into repopulating KCs in response to KC depletion, which is termed the “progenitor cell response”. Given our results show that repopulating KCs originate from non-monocytic hematopoietic progenitor cells in the BM, the progenitor cell response elicited by KC depletion is defined here as proliferation in BM, mobilization from BM into circulation, engraftment in the liver, and differentiation into KCs. Then, we sought to investigate that triggered by KC depletion, which type of non-monocytic hematopoietic progenitor cells proliferate in the BM, mobilize from BM into circulation, engraft in the liver, and differentiate into KCs.

For this purpose, we depleted KCs in C57BL/6 mice by intraperitoneal injection with 20 mg/kg of Clo and tracked the number of HSCs during KC repopulation. We found that the number of HSCs was dramatically increased at 48 hours post Clo injection (2 days before the start of KC repopulation), peaked at 96 hours, returned to a normal value at 240 hours (Fig. 4*A* and Fig. S5). Furthermore, we compared the percentage of 5-ethynyl-2’-deoxyuridine (Edu) positive HSCs from mice at 0 and 48 hours post Clo injection. We found that the percentage of Edu^+^ HSCs in mice from the 48 hours post Clo injection group was 60% greater than that in the 0 hours post Clo injection group (Fig. 4 *B* and *C*) [34.58 ± 5.19% vs. 10.40 ± 3.51%]. These findings indicated that following KC depletion, HSCs proliferated in the BM.

**Fig. 4.**
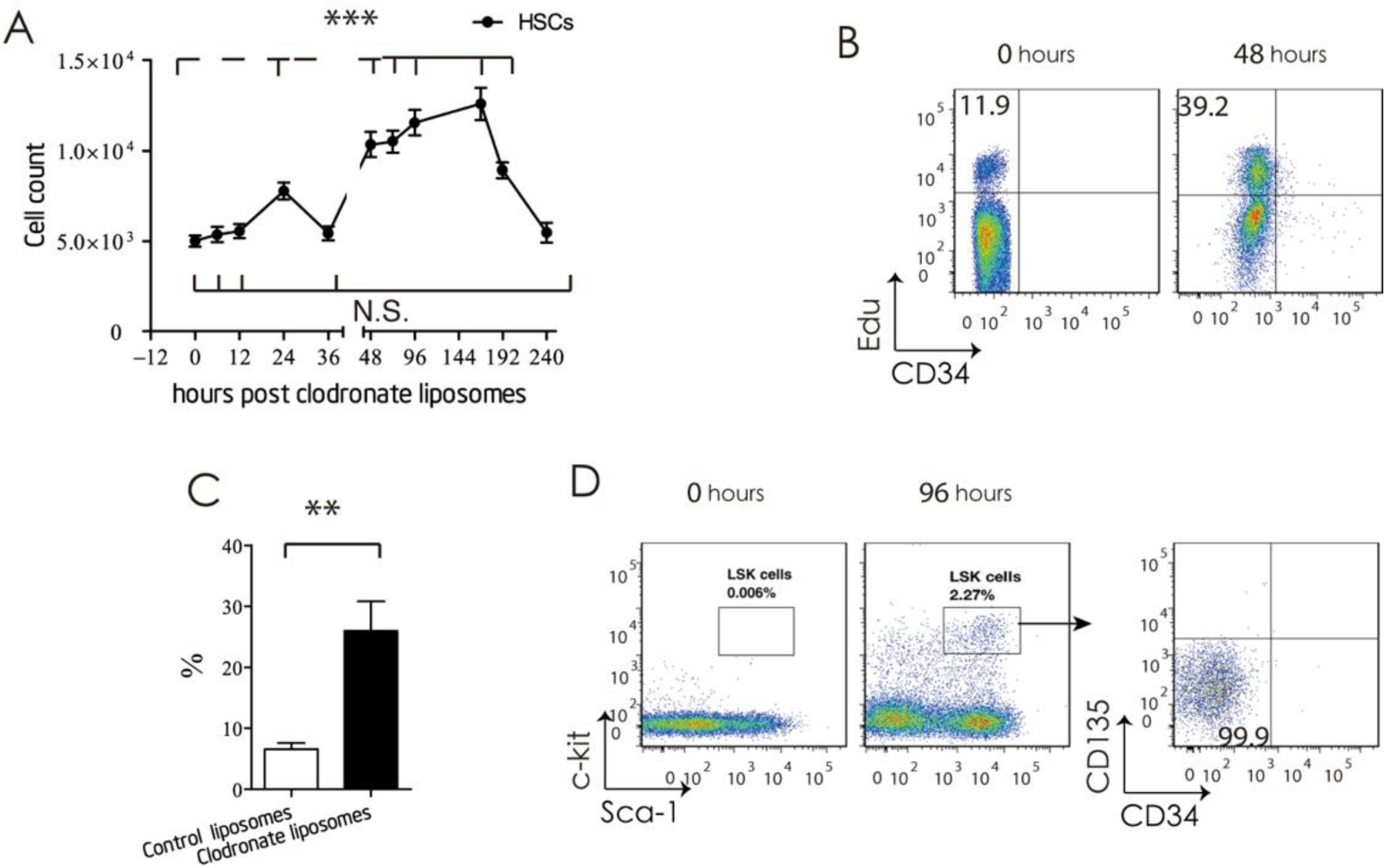
Hematopoietic stem cells proliferate in the bone marrow and mobilize into the blood, in response to Kupffer cell depletion. (*A*) Cell count of HSCs from mice i.p. with 20mg/kg clodronate-liposomes at indicated time point post injection. Values are the means ± SEM from 6 samples. \*\*\**P* < 0.001 between groups by ANOVA. (*B*) flow cytometric analysis of HSCs from mice i.p. with 20mg/kg clodronate-liposomes at indicated time point post injection (*C*) Percentage of 5-Ethynyl-2’-deoxyuridine (Edu) ^+^ HSCs from mice i.p. with 20mg/kg clodronate-liposomes at 48 hour post injection. Values are the means ± SEM from 6 samples. ** *P* < 0.01 between groups by *t*-test. (*D*) Flow cytometric analysis of blood Lin^eng^ cells from C57BL/6 mice at 4 day after control-liposomes or clodronate-liposomes injection.

Next, we performed time course analysis of HSC-specific markers on blood cells from mice received Clo injection. We found that a group of cells that express HSCs specific markers (Lin^neg^ Sca-1^+^ c-kit^+^, CD34^−^ CD135^−^) were detected in the blood for an observation period from 48 to 240 hours post Clo injection (Fig. 4*D* and Fig. S6), indicating that HSCs mobilized from the bone marrow into the blood during KC repopulation.

To investigate whether HSCs engraft in the liver and differentiate into KCs, we performed KC depletion in both B6GFP and C57BL/6 mice using intraperitoneal injection with 20mg/kg of Clo. Five days post-injection, we engrafted purified HSCs, multipotent progenitors (MPPs, defined as Lin ^neg^ Sca-1^+^ C-kit^+^ CD34^+^), or MOs from the BM of Clo treated B6GFP mice into different Clo treated C57/BL6 mice (Fig. 5*A*). Two days post-engraftment, we investigated the expression of HSC-specific markers Sca-1 and c-kit with flow cytometry on donor-origin NPCs isolate from HSC recipients. We found that Sca-1 and c-kit double-positive cells were detected in donor origin F4/80^−^ NPCs (Fig. 5*B*, and Fig. S7), indicating that transferred donor HSCs adoptively transferred into the liver of recipient. Then, 10 days post-engraftment, we analyzed the donor-origin marker GFP on F4/80^+^ KCs from HSC, MPP, and monocyte recipients. We found that 4.30 ± 1.36% of F4/80 ^+^ KCs were GFP positive in the HSC recipients, compared with less than 0.09 ± 0.07% in the MPPs recipients and 0.01 ± 0.005% in the monocyte recipients (Fig. 5 *C* and *D*), indicating transferred donor HSCs differentiated into KCs following engraftment in the liver of recipients. Finally, 90 days post engraftment, the donor chimerism of repopulating KCs in recipients were remained unchanged (Fig. 5*E*), indicating that the repopulating KCs can exist over the long term. All together, these data indicated that HSCs act as progenitor cells in response to KC depletion, including proliferation in the BM, mobilization into the blood, engraftment in the liver, and differentiation into KCs.

**Fig. 5.**
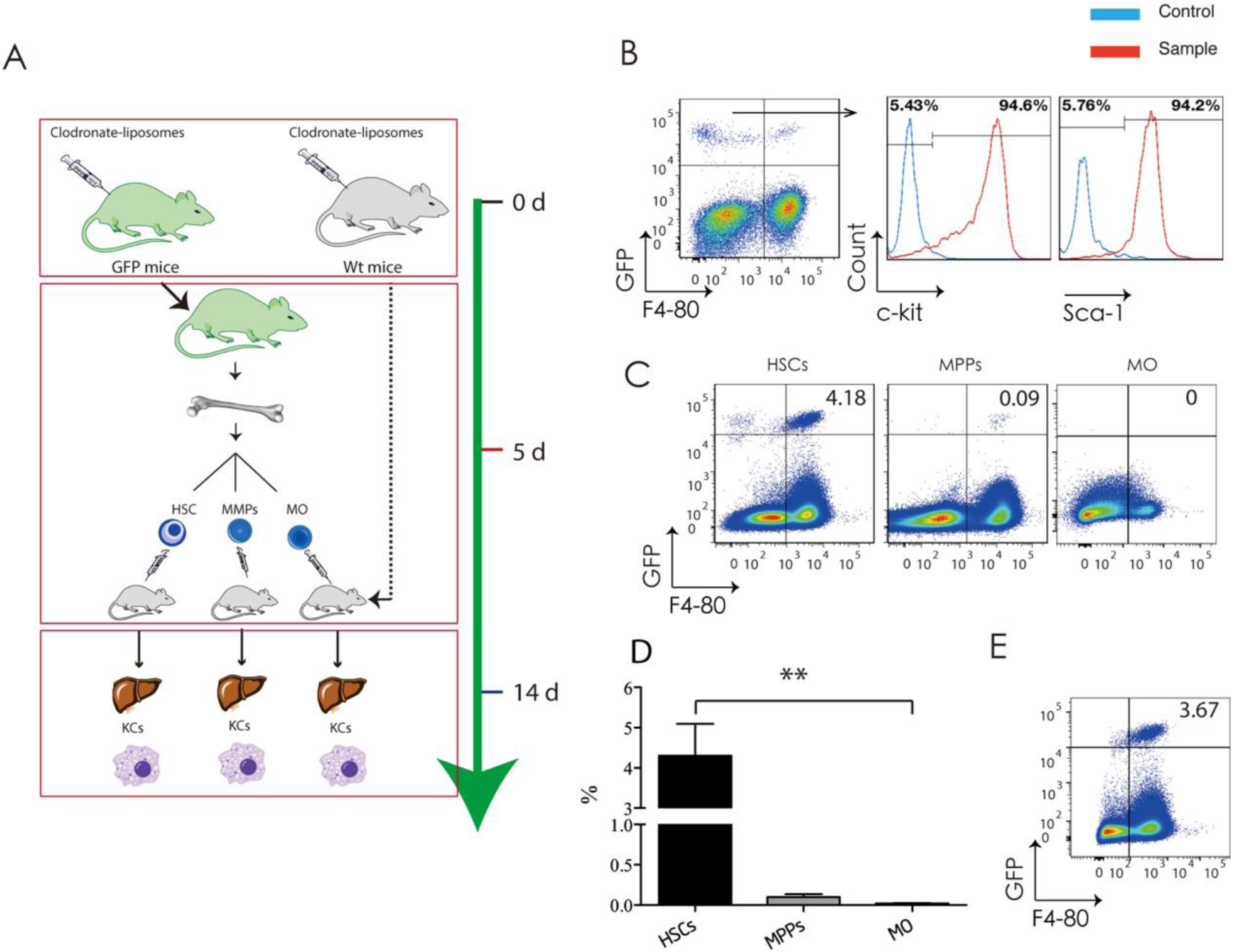
Hematopoietic stem cells adoptively transferr into the liver and differentiate into Kupffer cells, in response to Kupffer cell depletion. (*A*) Experimental schedule for adaptive transfer. (*B*)Flow cytometric analysis of liver non-parenchymal cells (NPCs) from KC-depleted or purified GFP^+^ HSCs-engrafted C57BL/6 mice at 2 day post-engraftment. (*C*) Flow cytometric analysis of NPCs from KC-depleted and purified GFP^+^ HSCs-engrafted, KC-depleted and purified GFP^+^MPP engrafted, or KC-depleted and purified GFP^+^ MOs (MOs) engrafted C57BL/6 mice at 10-day post-engraftment. (*D*) Percent of GFP^+^ KCs analyzed in *C*. Values are the means ± SEM from 6 samples. \*\*\**P* < 0.001 between groups by ANOVA. (*E*) Flow cytometric analysis of liver NPCs from KC-depleted or purified GFP^+^ HSCs-engrafted C57BL/6 mice at 90 day post-engraftment.

### Fate-mapping confirmed repopulating KCs originate directly from HSCs

Finally, we used a genetic inducible fate-mapping approach to confirm that repopulating KCs originate directly from HSCs, in a mouse model of chronic liver necro-inflammation induced by CCl_4_, in which resident KCs were depleted while the circulating MOs were not. The depletion of KCs was confirmed by tracking the number of GFP-resident KCs in the liver of non-myeloablative HSC chimeras treated with repeated CCl_4_. We found that the number of resident KCs decreased during repeated CCl_4_ treatment, and remained at the reduced level for the observation period from 0-d till 90-d after cessation of CCl_4_ treatment (Fig. 6 *A - C*). We also found a small amount of GFP^+^ macrophages in the liver of HSC-chimeras, for the observation period from 14-d till 90-d after cessation of CCl_4_ treatment, indicating that these cells were resident KCs cells rather than passenger inflammatory macrophages (Fig. 6 *A - C*). In short, these results suggested that during chronic liver inflammation induced by repeated CCl_4_ treatment, a small portion of embryonic derived KCs were depleted and replenished by hematopoietic progenitors.

**Fig. 6.**
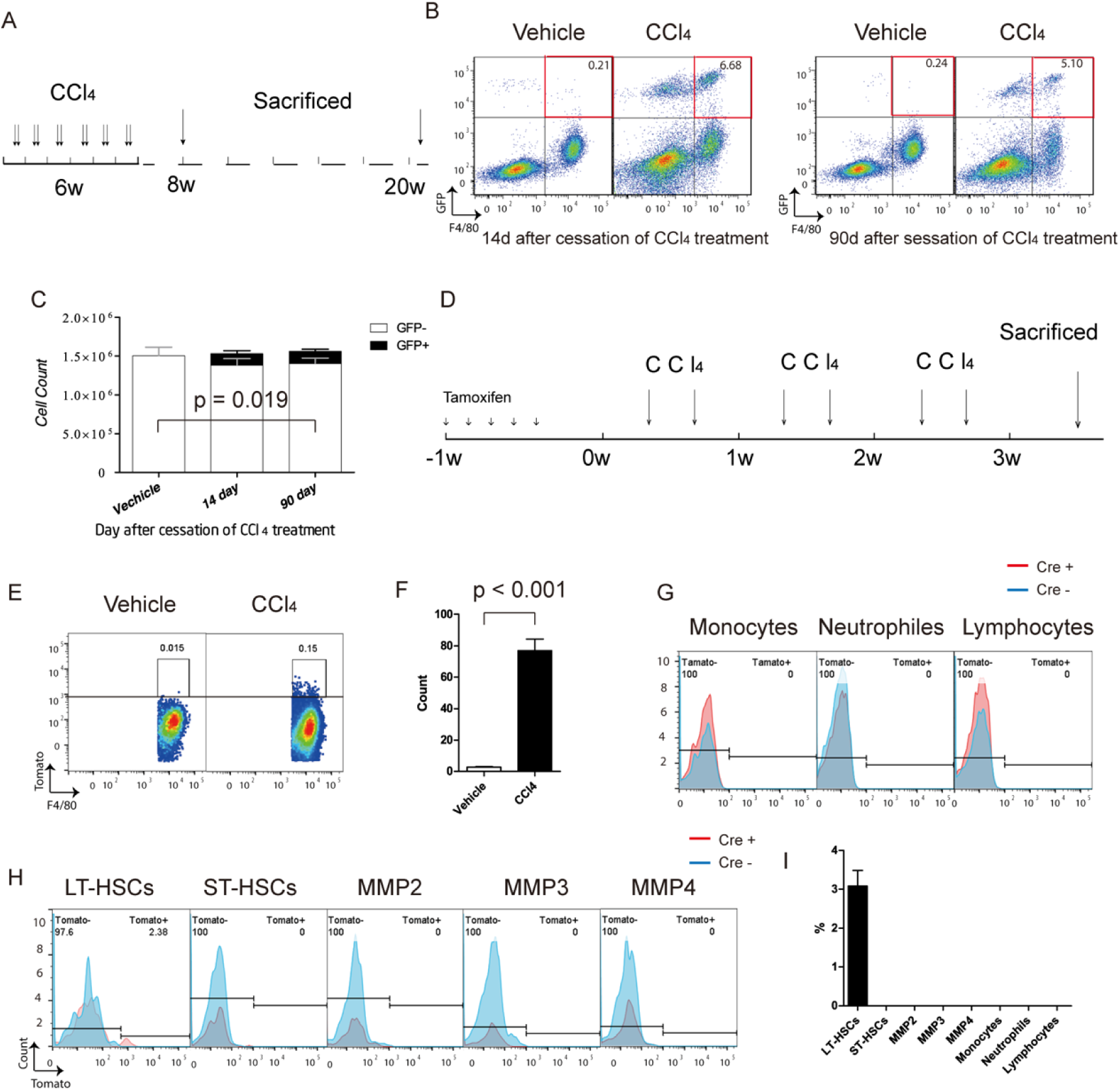
Repopulating KCs originate directly from HSCs, in context of CCl_4_ induced chronic liver inflammation. (*A*) Experimental schedule for counting the number of embryonic derived Kupffer cells (*B*) Flow cytometric analysis of liver non-parenchymal cells (NPCs) from repeated CCl_4_ or vehicle treatment purified GFP^+^ HSC-chimeric Kit^w^/Kit^wv^ mice at indicated time point. (*C*) Cell counts of GFP-embryonic derived KCs from HSC-chimeric Kit^w^/Kit^wv^ mice treated with repeated vehicle or CCl_4_ at 14 and 90 day after last dose of CCl_4_. Values are the means ± SEM from 4 samples analysis by ANOVA. (*D*) Experimental schedule for tracing the fate of HSCs during Kupffer cell repopulation in context of chronic liver inflammation induced by repeated CCl_4_. (*E*) Flow cytometric analysis of liver KCs from pulsed Fgd5-Cre ERT2/ stop-tdTomato-Cas9 mice received repeated vehicle or CCl_4_ treatment at 3-W after first dose vehicle or CCl_4_, respectively. (*F*) Cell count of Tomato^+^ KCs analyzed in *E*. Values are the means ± SEM from 6 samples. *** *P* < 0.001 between groups by ANOVA. (*G*) (*H*) Flow cytometric analysis of hematopoietic stem & progenitors in BM, or peripheral blood leukocytes as indicated from pulsed Fgd5^Cre ERT2^/ stop^tdTomato-Cas9^ mice received repeated vehicle or CCl_4_ treatment at 3-W after first dose vehicle or CCl_4_, respectively. (*I*) Percent of Tomato^+^ KCs analyzed in *G* and *H*. Values are the means ± SEM from 6 samples. \*\*\**P* < 0.001 between groups by ANOVA.

To further confirm that repopulating KCs originate directly from HSCs, we employed a HSC specific fate-mapping system constructed by crossing Fgd5-Cre ^ERT2^ mice^24^ with stop^tdTomato-Cas9^ mice, in which HSCs rather than MOs are genetically labeled. The rationale for this approach is that in this HSC specific fate-mapping system, the time to reach equilibrium between labeling index of HSCs and their progeny is especially long because of the exceedingly long residence time of short-term HSCs (ST-HSCs) and of Multipotent progenitors (MPPs)^25^. And then, we traced the fate of HSCs during chronic liver inflammation induced by repeated CCl_4_ treatment. To labeled HSCs, Adult Fgd5^Cre ERT^; ROSA26 ^stop-tdTomato-Cas9^ mice were pulsed with 5 days of consecutive of tamoxifen administration. To deplete KCs, 3 days after the pulse the mice were repeatedly injected with CCl_4_. To confirm that repopulating KCs originate directly from HSCs, not from MOs, the labeling index of repopulating KCs, of hematopoietic stem & progenitor cells in the BM, and that of peripheral blood leukocytes were detected at 21 days after the initiation of CCl_4_ treatment. We found that besides HSCs (LT-HSCs) only repopulating KCs were genetically labeled, in the HSC specific fate-mapping system (Fig. 6 *D – I*). These results indicated that repopulating KCs originate directly from HSCs, not from MOs or from monocytic progenitors.

## Discussion

In this study, in context of selective KC-depletion induced by Clo or by repeated CCl_4_ treatment, we provide in vivo fate-mapping evidences that repopulating KCs originate directly from HSCs, rather than preexisting KCs or MOs.

The long-standing notion that KCs have the potential to proliferate is based largely on measuring DNA-synthesizing KCs that can be labeled with thymidine analogs, in the context of inflammation or granuloma formation11. However, it is important to note that these thymidine analogs can be taken by cells undergoing abortive mitosis and by cells repairing DNA^26^. That means incorporation of thymidine analogs does not always mean proliferation of KCs. Most importantly, these studies actually test cell potential instead of cell fate. In our study, the contribution of preexisting KCs to KC repopulation was excluded by directly tracing the fate of preexisting KCs, a widely used method to determine the extent to which putative progenitor cells contribute to tissue regeneration^27^. In line with this finding, we also demonstrated that all repopulating KCs were of hematopoietic origin in non-myeloablation HSC chimeras.

A recent lineage-tracing study reported that Ly6C^hi^ monocytes gave rise to repopulating KCs in liver-shielded BM-chimeras in which KCs depletion is triggered with diphtheria toxin (DT) administrations^8^. However, it is important to note that inflammatory response triggered by DT administration^28–30^ can result in the recruitment of MOs, which can further differentiate into inflammatory macrophages in the liver^31^. Perhaps, that is why the number of Ly6C^hi^ MOs increased in the initial stages of KC repopulation following DT depletion. For the same reason, in DT treated CCR2^−/-^KC-DTR recipients, CCR2 expressed donor Ly6C^hi^ MOs were recruited into the liver by MCP-1, and differentiated into inflammatory macrophages which is difficult to be distinguished from KCs.

In our study, to investigate on the repopulation of KCs, not the infiltration of MOs, we firstly employed a Clo induced selective KC-depletion approach that did not trigger liver inflammation. Secondly, we employed genetic labeling approaches to distinguish monocyte-derived inflammatory macrophages from resident KCs, in context of CCl_4_ induced chronic liver inflammation.

We depleted KCs by Clo injection, which is a widely used and well defined model for investigating the function and repopulation of KCs. Although, like the common downside of currently available models for macrophage depletion, Clo injection depleted a broad range of mononuclear phagocyte cell types including BM MPS cells and circulating monocytes^32^. However, in our study, a low dose of Clo was injected intraperitoneally (Clo are substantial taken up by KCs due to absorption through portal circulation^33^), which selectively deplete KCs but not BM MPS cells. These results indicated that the mobilization of HSCs is a response to KC depletion, but not a response to depletion of BM MPS cells^34^.

In our monocytic cell-specific genetic inducible fate-mapping system, a few repopulating KCs are labeled. One explanation for this finding is that the Cx3cr1 promoter is expressed in the monocytic-intermediate between HSCs and repopulating KCs during differentiation; and the residual Cre activity induces gene reconstitution in a few of monocytic intermediates, even after a 15-day wash-out period. This presumption is supported by a recent study reported that although KCs ceased to express the Cx3cr1 chemokine receptor, they obviously originated from Cx3cr1 expressing precursors^20^. The possibility of MOs contribute to KC repopulation were further excluded by the finding that repopulating KCs are labeled in a HSC-specific fate-mapping system.

In summary, using genetic inducible fate-mapping approaches, we provide strong in vivo evidences that repopulating KCs do not originate from preexisting KCs or from MOs, but instead originate directly from HSCs. Our findings may shed light on the divergent roles of KCs in liver homeostasis and diseases.

## Materials and methods

### Mice strains and procedures

Tg(Csf1r-Mer-iCre-Mer)1Jwp mice (Jax#019098), R26R-EYFP mice (Jax#006148), W/Wv mice (Jax#100410), mT/mG mice (Jax#007676), Cx3cr1^<tm2.1(cre/ERT2)Jung>/^J mice (Jax#020940), Fgd5^ZsGr.CreERT2^ mice (Jax#027789), B6 ACTb-EGFP mice (Jax#003291) were purchased from the Jackson Laboratory. Stop-Cas9 mic (#T002249) were purchased from NanJing Biomedical Research Institute of Nanjing University (China). Wildtype C57BL/6 mice were obtained from the Institute of Laboratory Animal Science Chinese Academy of Medical Science. CCR2-mice (Jax#004999) were kindly provided by Dr. Li Tang (Beijing Institute of Lifeomics). Unless otherwise stated, mice were used at 6-12 weeks of age. Experimental mice were age- and sex-matched.

The investigators were blinded to the genotype of the animals during the experimental procedure. All experiments included littermate controls. Embryonic development was estimated considering the day of vaginal plug formation as 0.5 days post-coitum (dpc). All mice were bred and maintained in specific pathogen-free facilities at the Beijing Friendship Hospital. All animal procedures performed in this study were approved by the Institutional Animal Care and Use Committee of Capital Medical University. Reagents were from Sigma-Aldrich (Poole, UK) unless otherwise specified.

PCR genotyping of FVB-Tg^(Csf1r-cre/Esr1*)1Jwp/^J, B6.129×1-Gt(ROSA)26Sor^tm1(EYFP)Cos/^J, WBB6F1/J-^Kitw/Kitw-v/^J, B6.129(Cg)-Gt(ROSA)26Sor^tm4(ACTB-tdTomato,-EGFP)Luo^, B6.129P2(C)-Cx3cr1^<tm2.1(cre/ERT2)Jung>/^J, C57BL/6N-Fgd5^tm3(cre/ERT2)Djr/^J, and C57BL/6-Tg^(CAG-EGFP)10sb/^J mice, B6-Gt(ROSA)26Sor^tm1(CAG-LSL-cas9,-tdTomato)/^Nju mice, and B6.129^S4-Ccr2tmi1lfc/^J mice was performed according to the manufacturer’s instructions.

### Pulse labelling of Csf1r^+^ progenitors in embryos, Cx3cr1^+^ monocytic cells and Fgd5^+^ HSCs in adults

For genetic cell labelling of Csf1r^+^ progenitors in embryos, mice embryos recombination was induced by single injection of 75 μg/g (body weight) of tamoxifen (Sigma, T-5648) into pregnant females. To counteract the mixed estrogen agonist effects of tamoxifen, which can result in late fetal abortions, progesterone (Sigma, P-3972) dissolved in sterile vegetable oil was added for IP injections into pregnant females.

For genetic cell labelling Cx3cr1^+^ monocytic cells and Fgd5^+^ HSCs in adults, adult-mice recombination was induced by a 5 days consecutive injection of 200 μg/g (body weight) of tamoxifen.

### Isolation of cells from the blood, bone marrow and liver

<u>Blood cells</u> were collected as previously described^35^ before analysis by flow cytometry. Briefly, each mouse was humanely restrained in a modified plastic tube, exposing one of the hind limbs. The hair was removed using electric clippers and a thin layer of petroleum jelly was applied to the skin. The saphenous vein was punctured using a sterile 4 mm lancet and blood was collected into a microvette tube containing 2 mg/ml EDTA.

<u>Bone marrow cells</u> were collected as previously described^36^ before analysis by flow cytometry. Briefly, sacrificed mice were immersed in 75% ethanol. The skin was clipped mid-back and removed from the lower part of the body. The tissue was removed from the legs with scissors and dissected away from the body. Each end of the bone was cut off, and, using a 27 g needle/1 ml syringe filled with PBS, the bone marrow was expelled from both ends of the bone with a jet of medium directed into a 15 ml cell culture dish. The cell suspension was filtered through a 70-µm filter mesh to remove any bone spicules or muscle and cell clumps.

<u>Non-parenchymal liver cells</u> were isolated as described previously^37^. In short, the liver was perfused with collagenase and incubated at 25°C for 10 min in DNAse I solution. After collagenase digestion was halted with 5 mM EDTA solution, the resulting single-cell suspension was subjected to velocity and density centrifugation in an iodixanol gradient (Axis-Shield, Oslo, Norway) to produce purified suspensions of non-parenchymal cells.

### Flow cytometry

Erythrocytes in the blood were lysed using FACS^lyse^ solution (BD Pharmingen San Diago, CA). The isolated cells were surface stained in FACS buffer (PBS w/o Ca^2+^ Mg^2+^ supplemented with 0.5% BSA and 5 mM EDTA) for 30 min on ice. Multi-parameter analysis and flow cytometric cell sorting were performed on a FACS Aria II (BD Biosciences San Jose, CA) and analyzed with FlowJo software (Tree Star, Inc., Ashland, OR, USA).

For absolute F4/80^+^ cell counts, total NPCs isolated from each mouse were stained and sorted separately, and the cell number was counted with flow cytometry during FACS.

Fluorochrome-conjugated mAbs specific to mouse F4/80 (clone BM8), CD115 (clone AFS98), Ly6C (HK 1.4), Ly6A/E (clone D7), CD117 (clone 2B8), CD135 (clone A2F 10), CD34 (clone RAM34), CD207 (clone 4C7), I-A/I-E (clone M5/114.15.2), and the corresponding isotype controls were purchased from BioLegend (San Diego, CA, USA.). CD3e (clone 145-2c 11), CD19 (clone 1D3), and a lineage cocktail with an isotype control (561317) and the Annexin:PE Apoptosis Detection Kit I were purchased from BD Pharmingen (San Diago, CA).

### Transplantation of HSCs without irradiation

HSC transplantation in non-irradiated Kit^W/Wv^ mice was performed as described previously. In brief, approximately 2000 HSCs (Lin^neg^ Sca-1+ c-kit+, CD34- CD135-) isolated from the bone marrow of 3-weeks-old B6GFP mice, which carry a constitutively active EGFP reporter allele, were injected into 16-weeks-old Kit^W/Wv^ mice. Recipients were analyzed 8 weeks after transplantation for donor/host chimaerism in bone marrow, blood and liver.

### Kupffer cell depletion with clodronate liposomes

1:1 PBS-diluted clodronate liposomes and control liposomes (FormuMax Scientific, Palo Alto CA, USA) were injected via the Intraperitoneal as 20mg/kg, or 10mg/kg.

### Experimental model of liver injury

#### Acute liver injury

Mice received 0.6 mL/kg body weight of CCl_4_ mixed with corn oil intraperitoneally and were sacrificed at the indicated time points.

#### Chronic liver injury

CCl_4_ was injected twice weekly for 6 weeks. Mice were sacrificed at the indicated time point after the last injection.

### Magnetic enrichment of lineage-cells from single cell suspension of bone marrow

Depletion of lineage-committed cells from single-cell suspensions of mouse bone marrow was performed using the EasySep™ Mouse Hematopoietic Progenitor Cell Isolation Kit according to the manufacturer’s instructions (StemCell Technologies, Vancouver, BC, Canada).

### Labeling DNA of proliferating cells with 5-ethynyl-2’-deoxyuridine in vivo

Incorporation of 5-ethynyl-2’-deoxyuridine (Edu) was measured using the Click-iT EdU flow cytometry assay kit according to the manufacturer’s instructions (Life technologies, Carlsbad, CA, USA). Briefly, Edu was dissolved in DMSO at 25.5 mg/ml and further diluted in PBS to 5 mg/ml. Mice were injected i.v. with 50 μg/g Edu 1 day prior to sacrifice. Controls received DMSO/PBS.

### Histopathological Examination

Each formaldehyde-fixed sample was embedded in paraffin, cut into 5 μm-thick sections and stained with hematoxylin-eosin (H-E) according to standard procedures. All slides were reviewed by the same pathologist.

### Cell transfer

Sorted populations isolated from the bone marrow of every clodronate-liposome-treated, B6GFP^+^ donor mouse at 5 days post clodronate-liposome injection (HSCs, 3.0 × 10^4^ cells per mouse; MPPs, 3.0 × 10^4^ cells per mouse and MO, 5.0 × 10^5^ cells per mouse) were injected intravenously into each clodronate-liposome-treated C57BL/6 mice at 5 days post clodronate-liposome injection.

### Statistical analysis

Result represent the mean ± s.e.m. unless otherwise indicated. Statistical significance was determined as indicated in figure legends. Statistical analyses were done with Prism GraphPad software v5.0, and the exact tests used are indicated within the appropriate text.

## Author Contributions

X. F. designed the study, performed most of the experiments, analyzed the data, and wrote the manuscript. XH. C., P. L., H. Y., D. Z. and HD. W. performed the fate-mapping experiments. WR. L., FY. C., and P. W. performed the PCR analysis. Y. J., JD. J., and FC. H. supervised the study, Y. J. designed most of the experiments and revised the manuscript. JD. J. helped revise the manuscript. SZ. Y. and B. L. helped design the fate-mapping experiments and B. L. revise the manuscript. All authors contributed to the manuscript.

## Acknowledgments

We are grateful to W. Liu and other members of the Beijing center excellence, Becton-Dickinson Biosciences for flow cytometry technical assistance. This work was partially supported by National Key R&D Program of China (Nos.2018YFA0507502, 2020YFE0202200), the National Natural Science Foundation of China (Nos. 81770581, 81570526 and 81770598), Beijing Science and Technology Project (Z161100002616036 and Z15111000160000), Open Project Program of the State Key Laboratory of Proteomics (SKLP-K201801 and SKLP-O201509), and the China Postdoctoral Science Foundation (2014M552634).

---

**Figure S1.**
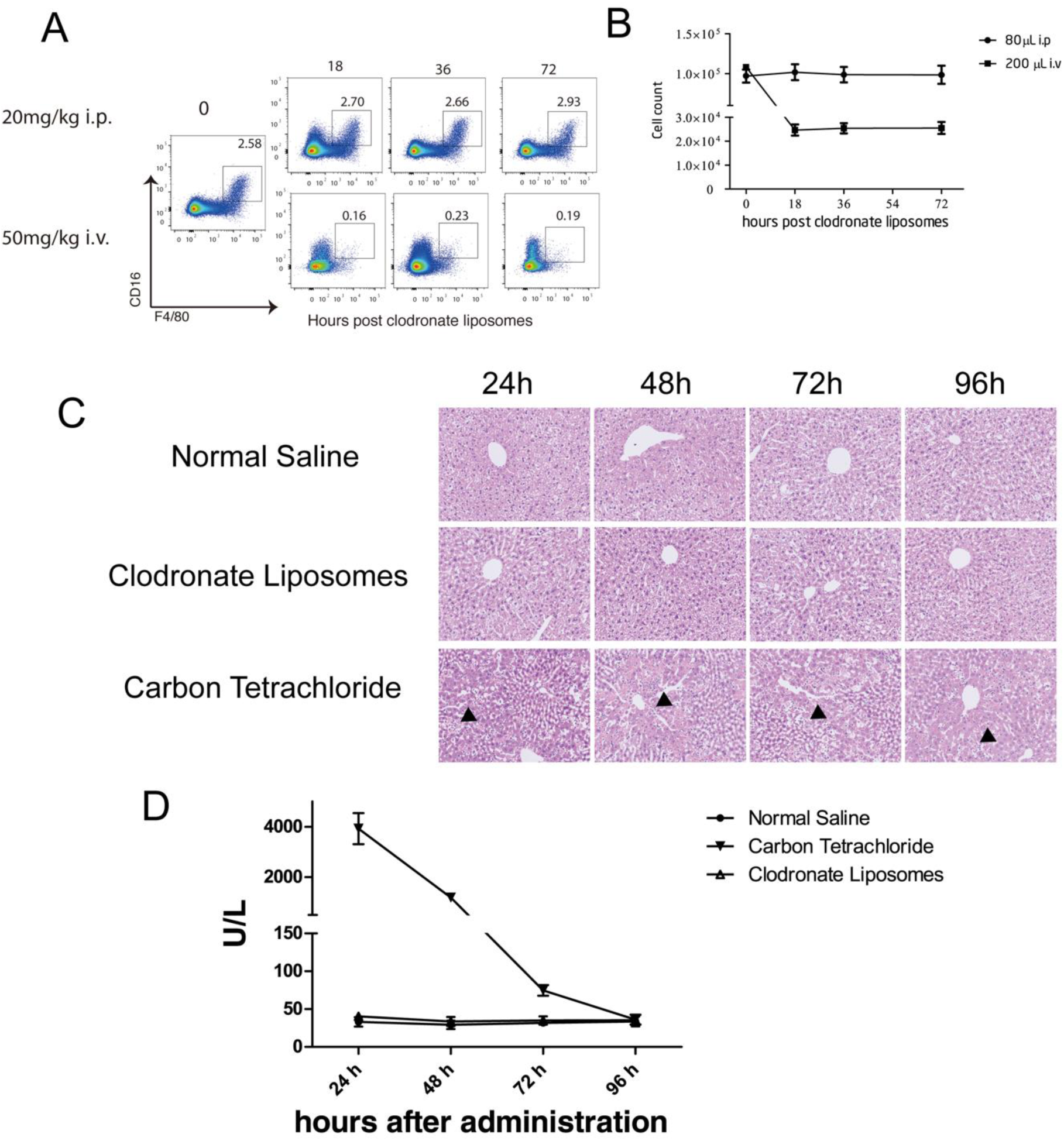
Intraperitoneal injection with 20mg/kg clodronate-liposomes did not deplete bone marrow macrophages, and did not trigger liver inflammation. (*A*) Flow cytometric analysis of BM MPS of C57/BL mice received indicated dose of Clo injection. (n = 5/group). (*B*) Cell count of BM MPS of C57/BL mice treated with indicated dowse of Clo analyzed in A. (*C*) Liver tissue from all normal-saline treatment mice at each time point revealed normal cellular architecture (n = 5**/**group). Liver tissue from single dose Carbon tetrachloride treatment group revealed some damage of liver cells, inflammatory cells infiltration, fatty changes and centrilobular necrosis (n = 5/group). Liver tissue from Clodronate liposomes group revealed no damage of liver cells and inflammatory cells infiltration (n = 5**/**group). (*D*) Serum alanine aminotransferase of mice from Carbon Tetrachloride group were significantly increased at 24 hours post treatment, and returned to normal level at 96 hours (n = 5**/**group). In contrast, serum alanine aminotransferase of mice from Clodronate Liposomes group and Normal Saline group were remained unchanged, at the mean time.

**Figure S2.**
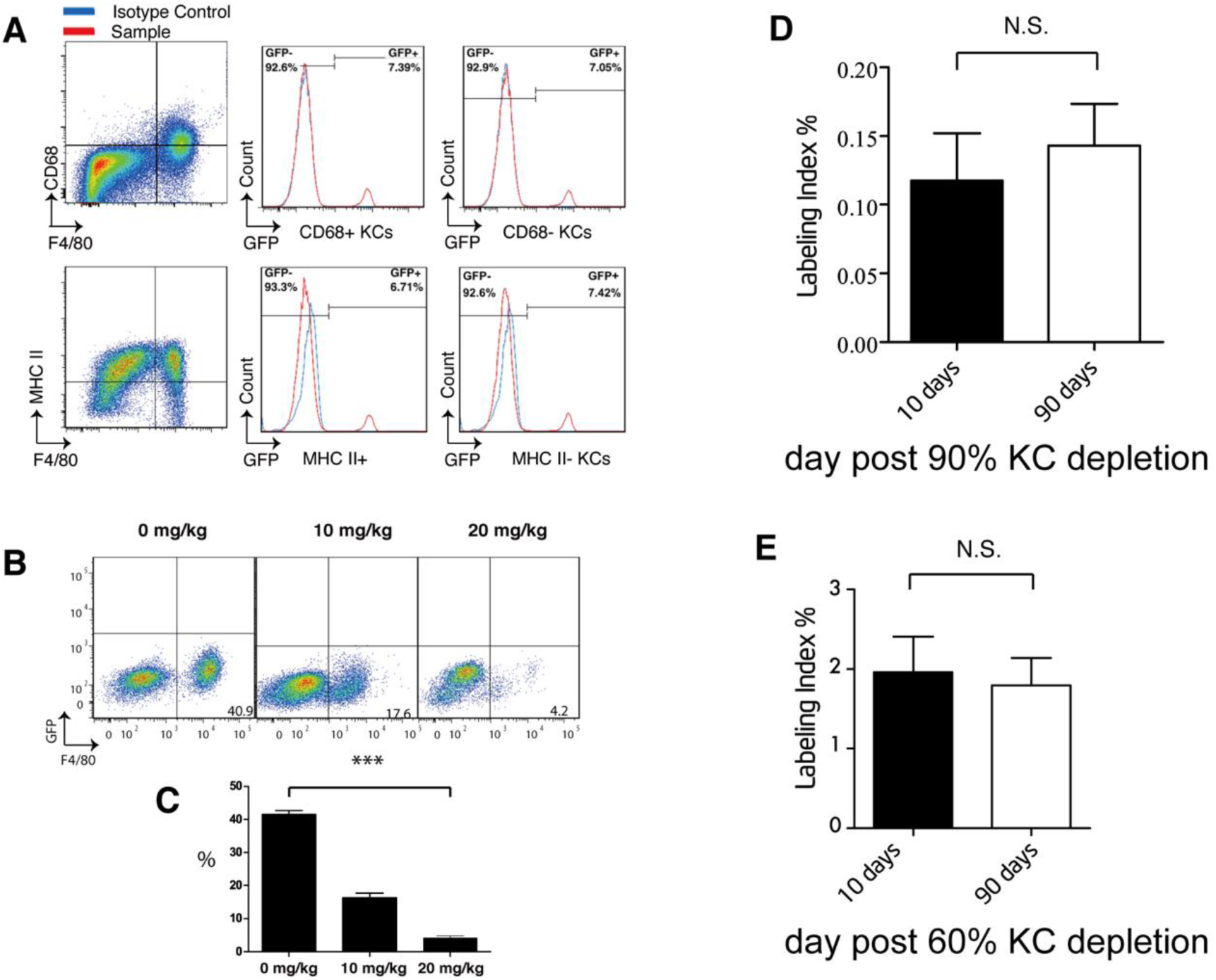
Analysis of Kupffer cells from C57B/L6 mice following intraperitoneal injection with 20mg/kg clodronate-liposomes. (*A*) GFP expression on CD68^+^ and CD68^−^ KCs from E8.5 pulsed Cre mice at 8 weeks after birth. (*B*) Flow-cytometric analysis of KCs from C57B/L 6 mice 24 hours after being treated with intraperitoneal injection of clodronate-liposomes of indicated dose (n = 6/group). (*C*) Percentage of KCs from C57B/L 6 mice treated with intraperitoneal injection of clodronate-liposomes of indicated dose analyzed in *B*. (*D*) Labeling index of KCs from E8.5 pulsed Csf1r^Wt^; Rosa^mT/mG^ mice at indicated time point post intraperitoneal injection with 20mg/kg Clo. N.S No significant difference. (*E*) Labeling index of KCs from E8.5 pulsed Csf1r^Wt^; Rosa^mT/mG^ mice at indicated time point post intraperitoneal injection with 10mg/kg Clo. N.S No significant difference.

**Figure S3.**
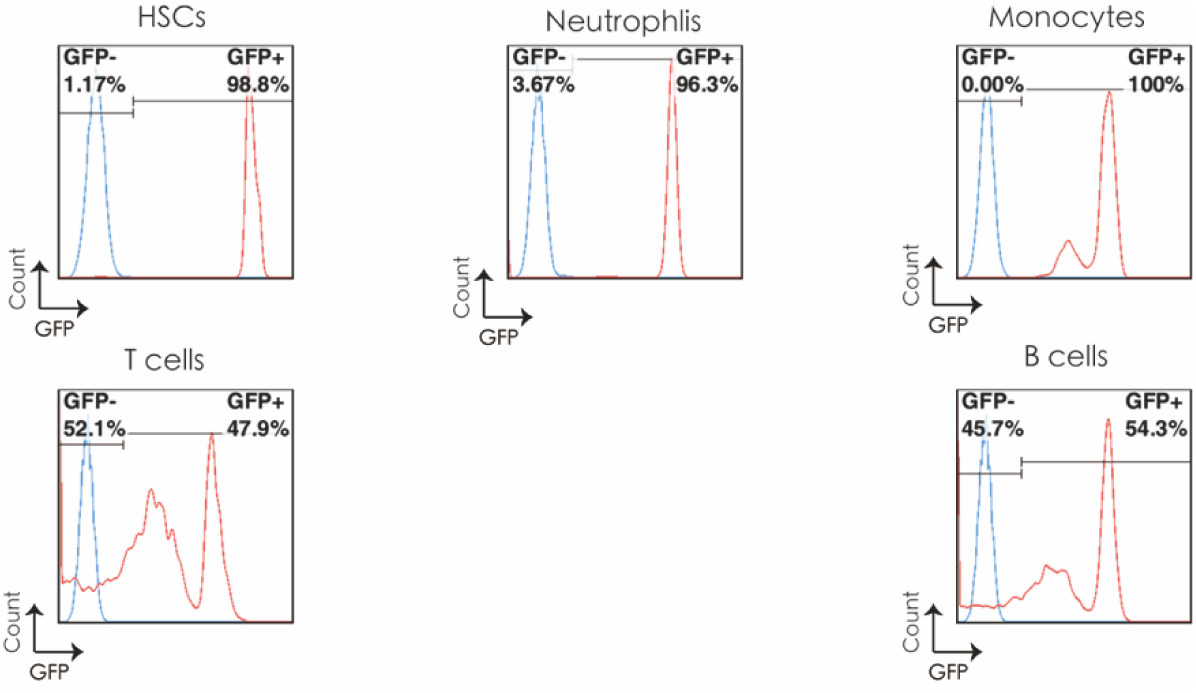
Flow cytometric analysis of GFP expression of hematopoietic stem cells and blood leukocytes within purified GFP^+^ HSC-chimeric Kit^w^/Kit^wv^ mice.

**Figure S4.**
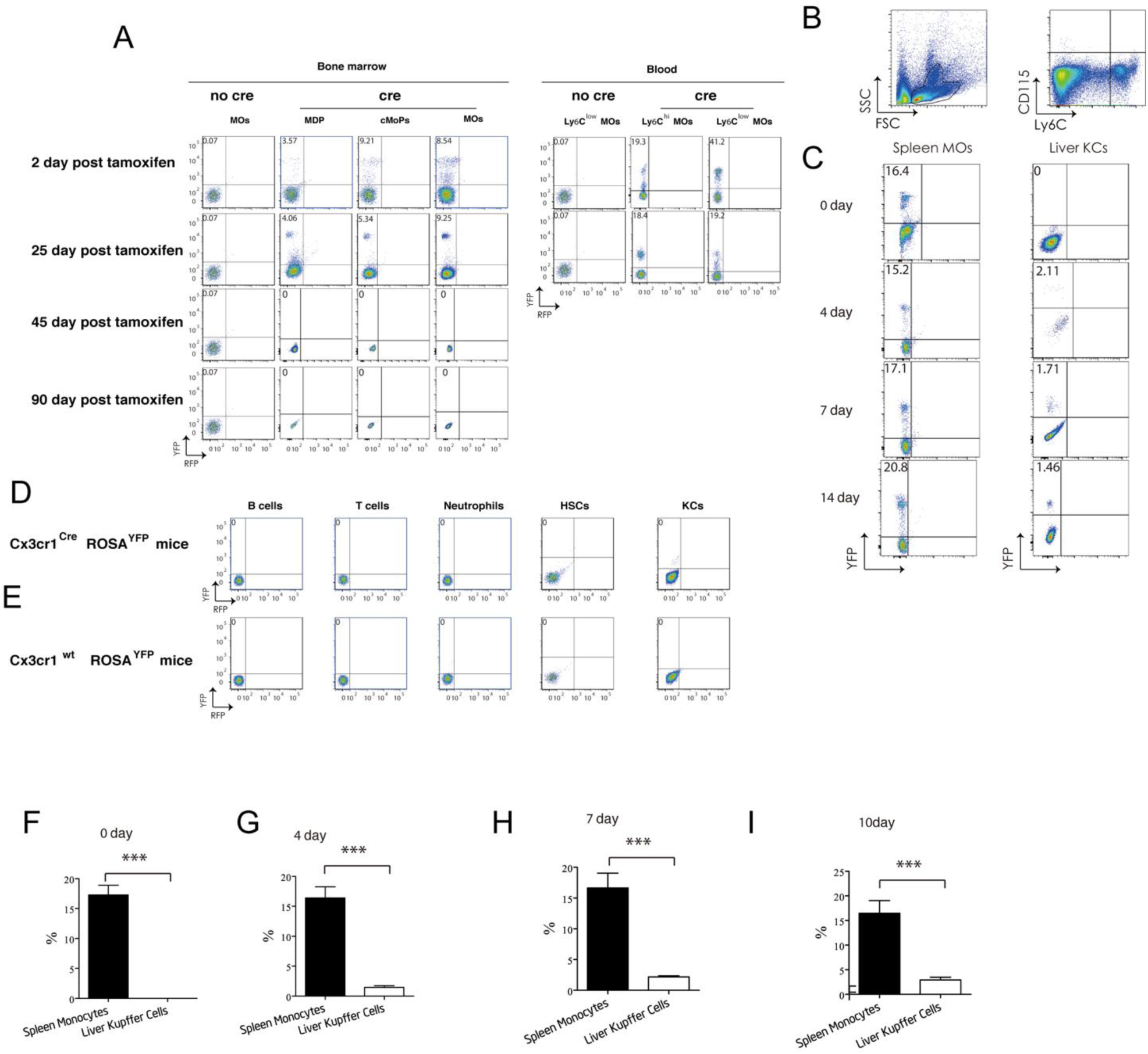
Flow cytometric analysis of YFP expression of indicated cells within adult pulsed Cx3cr1^CreERT2^; Rosa^YFP^ mice (Cre) or Cx3cr1^wt^; Rosa^YFP^ mice (No cre). (*A*) Flow-cytometric analysis of bone marrow monocytic cells and blood MO from adult pulsed Cx3cr1^wt^; Rosa^YFP^ or Cx3cr1^CreERT2^; Rosa^YFP^ mice at indicated time point post pulse (n = 5/group). (*B*) Gating strategy of intra-splenic MO. Dot plots are gated on viable single splenic cells. Intra-splenic MO are defined as Ly6C^+^ cells. (*C*) Flow cytometric analysis of YFP expression on intra-splenic MO and KCs within the same adult pulsed Csf1r^MeriCreMer^; Rosa^YFP^ mice at indicated time point post intraperitoneal injection of 20mg/kg Clo (n = 4/group). (*D*) Flow-cytometric analysis of blood leukocytes and KCs from adult pulsed Csf1r^MeriCreMer^; Rosa^YFP^ mice at 25-day post pulse (n = 5/group). (*E*) Flow-cytometric analysis of blood leukocytes and KCs from adult pulsed Csf1r^wt^; Rosa^YFP^ mice at 25-days post pulse (n = 5/group). (*F*), (*G*), (*H*), (*I*) Labeling index of intra-splenic MO and KCs at indicated time point post intraperitoneal injection of 20mg/kg Clo, analyzed in (*C*). *** *P* < 0.001.

**Figure S5.**
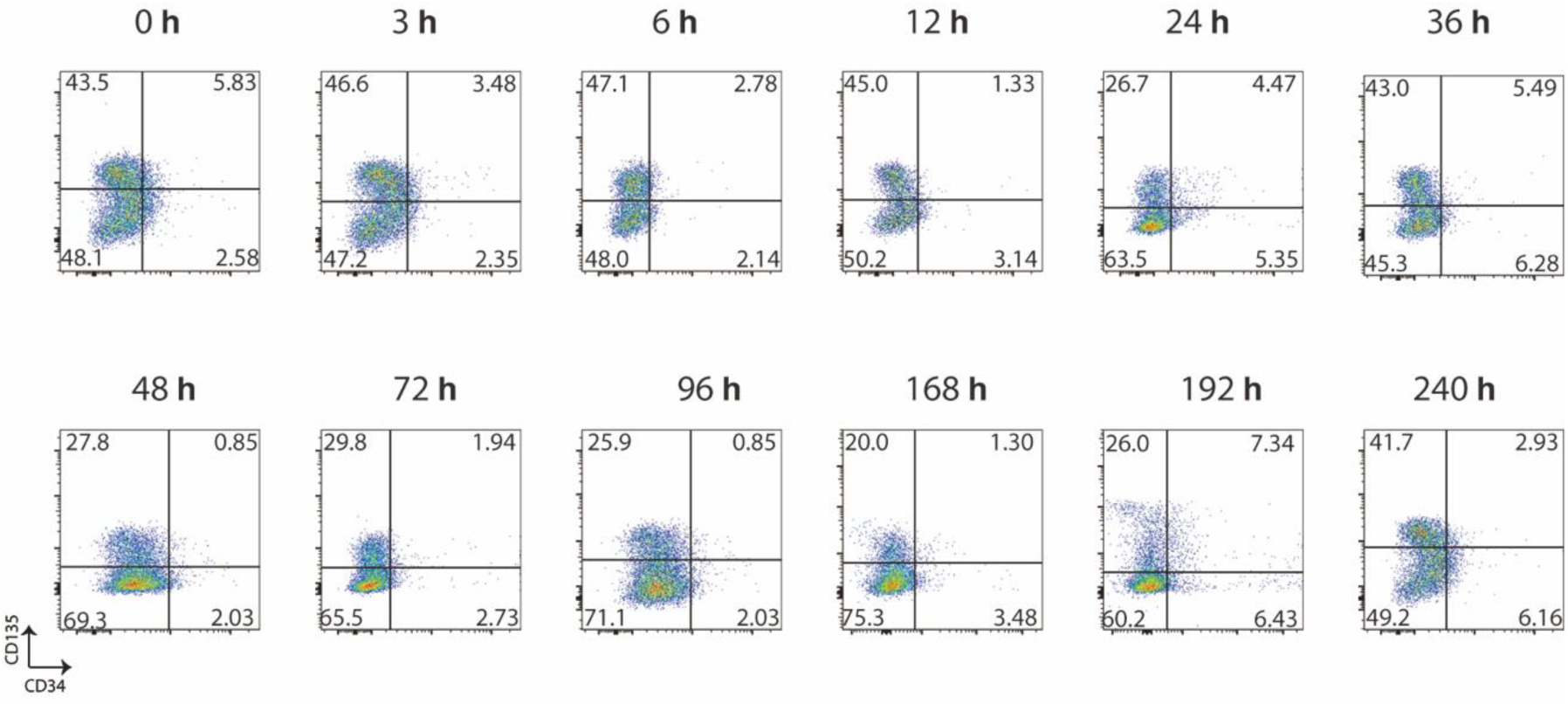
Flow-cytometric analysis of HSCs from C57B/L 6 mice at indicated time point post intraperitoneal injection with 20mg/kg Clo (n = 5/group).

**Figure S6.**
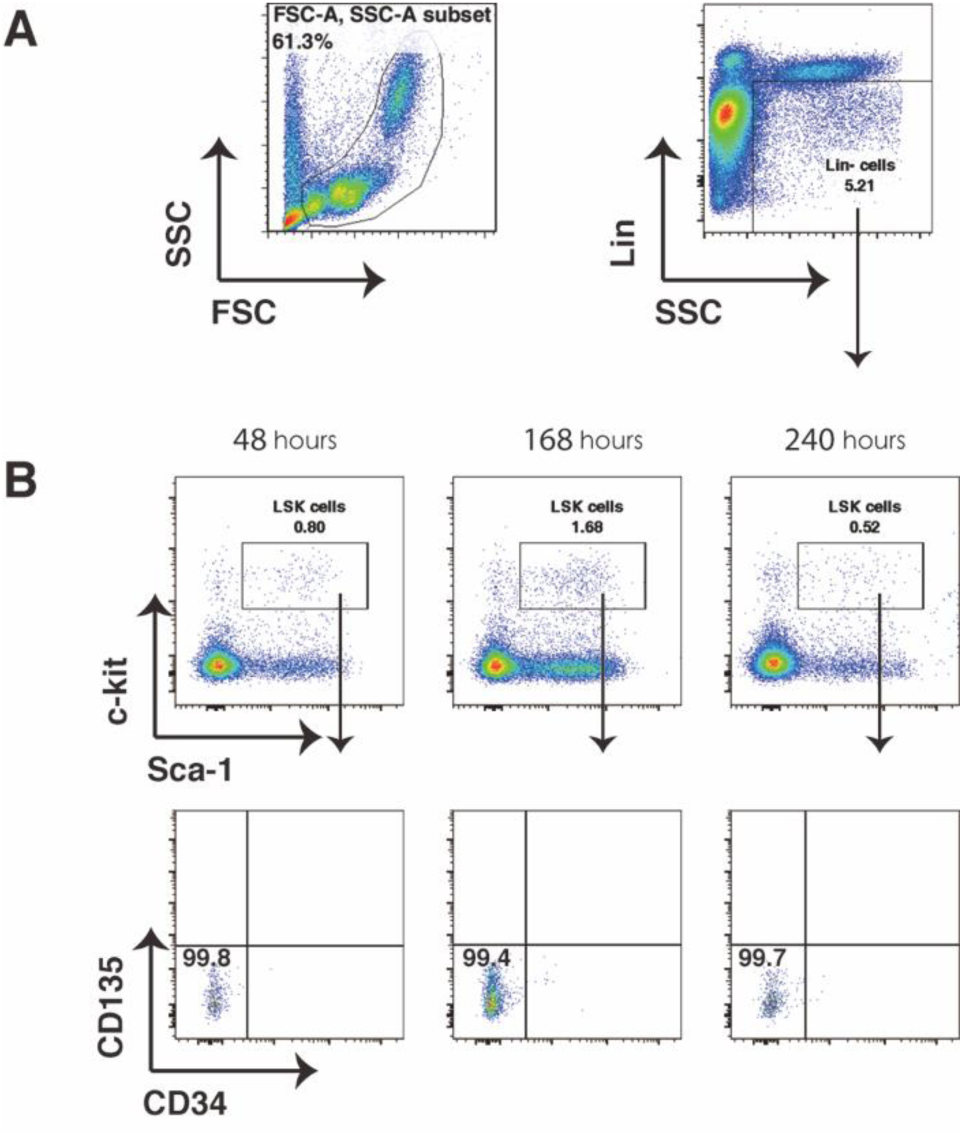
Flow-cytometric analysis of blood HSCs from C57B/L 6 mice at indicated time point post intraperitoneal injection with 20mg/kg Clo (n = 5/group). HSCs were defined as Lin^neg^ /Sca-1^+^ /c-kit^+^ /CD135^−^ /CD34^−^ cells.

**Figure S7.**
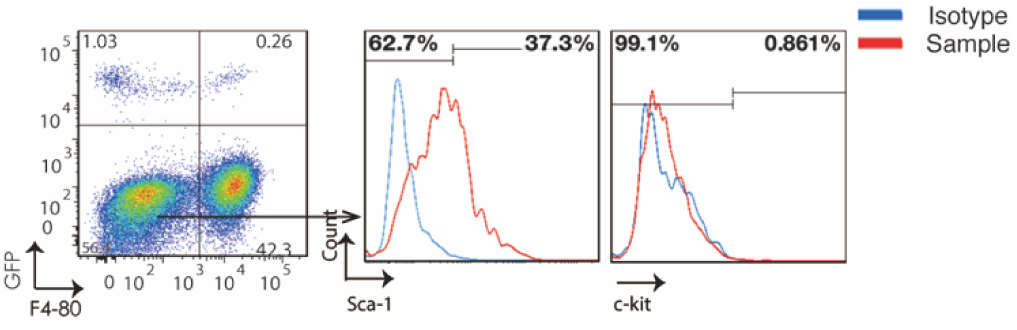
Flow cytometric analysis of GFP-liver non-parenchymal cells in KC-depleted mice revived GFP^+^ HSCs engraftment.

